# Transcription factor activity and nucleosome organisation in mitosis

**DOI:** 10.1101/392241

**Authors:** Nicola Festuccia, Nick Owens, Thaleia Papadopoulou, Inma Gonzalez, Alexandra Tachtsidi, Sandrine Vandoermel-Pournin, Elena Gallego, Nancy Gutierrez, Agnès Dubois, Michel Cohen-Tannoudj, Pablo Navarro

## Abstract

Mitotic bookmarking transcription factors (BFs) maintain the capacity to bind to their targets during mitosis, despite major rearrangements of the chromatin. While they were thought to propagate gene regulatory information through mitosis by statically occupying their DNA targets, it has recently become clear that BFs are highly dynamic in mitotic cells. This represents both a technical and a conceptual challenge to study and understand the function of BFs: first, formaldehyde has been suggested to be unable to efficiently capture these transient interactions, leading to profound contradictions in the literature; second, if BFs are not permanently bound to their targets during mitosis, it becomes unclear how they convey regulatory information to daughter cells. Here, comparing formaldehyde to alternative fixatives we clarify the nature of the chromosomal association of previously proposed BFs in embryonic stem cells: while Esrrb can be considered as a canonical BF that binds at selected regulatory regions in mitosis, Sox2 and Oct4 establish DNA sequence independent interactions with the mitotic chromosomes, either throughout the chromosomal arms (Sox2) or at pericentromeric regions (Oct4). Moreover, we show that ordered nucleosomal arrays are retained during mitosis at Esrrb book-marked sites, whereas regions losing transcription factor binding display a profound loss of order. By maintaining nucleosome positioning during mitosis, Esrrb might ensure the rapid post-mitotic re-establishment of functional regulatory complexes at selected enhancers and promoters. Our results provide a mechanistic framework that reconciles dynamic mitotic binding with the transmission of gene regulatory information across cell division.

## Introduction

During mitosis, the chromatin is drastically condensed and reconfigured to enable the equitable partition of the genetic material between the two daughter cells **(Ma et al. 2015)**. This leads to a strong decrease in transcriptional activity and to the general reduction of transcription factor (TF) binding throughout the genome. Loss of TF binding is further accentuated by the stereotypical phosphorylation of many regulators during mitosis, leading to an intrinsic reduction of their ability to bind DNA. This is particularly well illustrated by the systematic phosphorylation of C2H2 zinc finger TFs such as YY1 **(Rizkallah et al. 2011; Rizkallah and Hurt 2009)**, but has also been observed for other TFs such as Oct4 and Sox2 **(Shin et al. 2016; Qi et al. 2016)**. Moreover, the breakdown of the nuclear envelope, and the consequent increase of the volume that TFs can freely explore, leads to a decrease of TF concentration. This process naturally inhibits the ability of TFs to scan DNA for their binding motifs. Therefore, many processes occur simultaneously to temporarily halt gene regulation and transcription during mitosis. The mechanisms by which daughter cells accurately re-establish an environment permissive for efficient transcriptional activation early in interphase remain unknown **(de Castro et al. 2016)**. One potential mechanism is known as mitotic bookmarking: some TFs have the ability to interact with their DNA binding sites during cell division. These TFs, known as mitotic bookmarking factors (BFs), are believed to directly convey gene regulatory information from mother to daughter cells, as illustrated by Gata1 **(Kadauke et al. 2012)**, FoxA1 **(Caravaca et al. 2013)** and Esrrb **(Festuccia et al. 2016)**. Nonetheless, the molecular mechanisms underpinning this function remain to be elucidated **(Festuccia et al. 2017)**. BFs are highly dynamic during mitosis and often exhibit reduced residence times on the chromatin. Therefore, the function of BFs is not simply mediated by their stable retention at enhancers and promoters. Instead, their transient binding activity may preserve specific chromatin features at book-marked sites. These features would represent the inherited properties driving and accelerating the reassembly of functional regulatory complexes early in the following interphase. Remarkably, even though the chromatin is highly condensed during mitosis, gene regulatory elements remain globally accessible **(Hsiung et al. 2015)**. This is particularly true at active promoters, perhaps reflecting their low but nevertheless significant mitotic activity, as recently reported **(Palozola et al. 2017)**. Enhancers, in contrast, show more variable degrees of chromatin accessibility. Yet, mitotic chromatin accessibility does not seem to correlate with mitotic binding, at least in the case of bookmarking by Gata1 in erythroblasts **(Kadauke et al. 2012)**. Moreover, the maintenance of chromatin accessibility does not preclude the possibility that nucleosome positioning in mitotic cells is highly modified, as previously suggested **(Kelly et al. 2010)**. Hence, further studies are required to clarify whether regulatory elements do indeed maintain a local chromatin architecture compatible with TF binding in mitotic cells, and how mitotic bookmarking correlates with and ultimately drives nucleosome organisation. An essential condition to understand mitotic book-marking processes is to accurately identify BFs and their mitotic binding sites. However, this has remained a difficult task because, as reported nearly 15 years ago **(Pallier et al. 2003)**, the most commonly used cross-linker, formaldehyde, leads to the artificial depletion of TFs from mitotic chromosomes **(Pallier et al. 2003; Teves et al. 2016)**. To circumvent this problem mitotic bookmarking activity has been explored using live imaging of tagged TFs. Even so, whether the global chromatin association of certain TFs detected by microscopy reflects the sum of site-specific interactions remains to be demonstrated. Diverse modes of binding, others than those involving base-specific interactions, may be responsible for the macroscopic decoration of the chromosomes by TFs, as we proposed earlier **(Festuccia et al. 2017)** and was clearly demonstrated for FoxA1 **(Caravaca et al. 2013)**. These interactions with the chromatin, or with other constituents of mitotic chromosomes, might be extremely transient and not easily captured by formaldehyde. In support of this distinction between global and site-specific interactions, several TFs have been efficiently captured at their mitotic binding sites using formaldehyde **(Festuccia et al. 2017)**, despite its seeming incapacity to cross-link TFs on mitotic chromosomes. Yet, it remains to be proven whether formaldehyde generally fails in capturing transient DNA-specific interactions, leading to the loss of enrichment of BFs on the chromosomes, or whether the interactions sustaining the global retention of BFs are distinct from those involved in TF binding to DNA. This does not only represent an important technical question; rather, it directly interrogates the nature and, hence, the function, of the interactions established between TFs and mitotic chromosomes: while global, dynamic and DNA sequence independent interactions may increase the concentration of TFs in the vicinity of DNA, possibly facilitating the re-establishment of binding in the following interphase, authentic mitotic bookmarking of promoters and enhancers may confer specificity to these processes and provide robustness to the post-mitotic resuscitation of gene regulatory networks.

In this manuscript, we combine (para-)Formaldehyde (PFA) with alternative fixatives that are able to capture hyperdynamic protein-protein interactions, such as Disuccinimidyl Glutarate (DSG; **Tian et al. 2012)**, to study the association of TFs with mitotic chromatin. We report that DSG, in contrast to PFA, preserves the global interactions of two TFs, Esrrb and Sox2, previously proposed to display strong mitotic bookmarking activity in embryonic stem (ES) cells **(Deluz et al. 2016; Festuccia et al. 2016; Teves et al. 2016; Liu et al. 2017)**. Nonetheless, irrespective of the fixative used, DNA sequence-specific interactions can be detected for Esrrb at thousands of target sites, but not for Sox2. Moreover, imaging after DSG fixation unmasks a particular behaviour of another TF previously suggested to act as a BF, Oct4: both in interphase and in mitosis, Oct4 is focally enriched on pericentric heterochromatin. In mitosis, Oct4 is also largely excluded from the chromatid arms and, similarly to what we observe for Sox2, loses its ability to engage in site-specific interactions. These observations strongly argue against the idea that the global decoration of the mitotic chromosomes can be taken as an indication of a sequence-specific bookmarking activity. Integrating our genome-wide localisation studies with additional assays interrogating chromatin accessibility, nucleosome positioning and stability (Table S1), we conclude that mitotic bookmarking, particularly by Esrrb, is strongly associated with the preservation of an interphase-like nucleosomal organisation. At bookmarked sites, both in interphase and mitosis, Esrrb motifs lie at the centre of nucleosome depleted regions surrounded by phased arrays of nucleosomes. At loci losing TF binding, the mitotic nucleosomes are vastly reorganised and display increased fragility as compared to interphase. We conclude that TFs display a range of behaviours in mitotic cells that can be captured with distinct fixatives. This variability reflects the ability of some TFs, like Esrrb, but not of all factors enriched on mitotic chromosomes, to bind at specific regulatory elements efficiently and to maintain a nucleosomal organisation compatible with the rapid re-establishment of regulatory complexes in interphase.

## Results

### The global localisation of TFs to mitotic chromosomes is preserved upon DSG fixation

Several TFs have been shown to seemingly coat the mitotic chromosomes when fusion proteins with fluorescent proteins or tags are used in live imaging approaches **(Festuccia et al. 2017)**. This is the case of Esrrb **(Festuccia et al. 2016)**, which we previously showed to decorate mitotic ES cell chromatin using GFP (Fig. 1A), Tdtomato and Snap-tag fusions. However, upon Formaldehyde fixation, several TFs capable of coating the mitotic chromosomes, seem to be globally delocalised and crosslinked outside of the chromosomes **(Pallier et al. 2003; Teves et al. 2016)**. We first aimed to test whether this is also the case for Esrrb. As expected, we observed a clear depletion on the mitotic chromosomes (Fig. 1B), which were identified by DAPI (Fig. S1) and Ki-67 staining (Fig. 1B), a protein enriched on their periphery **(Booth and Earnshaw 2017)**. We therefore aimed at identifying alternative cross-linking agents that would preserve the chromosomal enrichment of Esrrb. Among the different reagents and protocols that we tested, we found two that clearly allow to visualise Esrrb coating of the mitotic chromosomes. First, the homobifunctional crosslinker DSG (Fig. 1B), which has been used to capture hyper-dynamic protein-protein interactions due to its capacity to establish amide bonds via two NHS-ester groups **(Tian et al. 2012)**. Both followed by PFA post-fixation (Fig. 1 and Fig. S1) or not (not shown), the staining patterns were found identical. Second, Glyoxal (Fig. S1), a small bifunctional aldehyde that has been recently rediscovered for its use in fluorescent microscopy **(Richter et al. 2018)**. As a control, we stained ES cells for Nanog (Fig. 1A), a TF that is excluded from the mitotic chromatin **(Festuccia et al. 2016)**. Upon DSG or Glyoxal fixation, we did not observe retention of Nanog on mitotic chromosomes (Fig. 1B and Fig. S1), indicating that these two cross-linkers do not induce aspecific aggregation on the chromosomes. We also tested whether DSG would allow us to visualise the global chromosomal retention of Esrrb in vivo. We have shown before that upon microinjection of Esrrb-Tdtomato mRNA into mouse embryos, the produced fluorescent fusion proteins decorate the mitotic chromatin **(Festuccia et al. 2016)**. Accordingly, when we fixed mouse blastocysts with DSG we could observe mitotic figures with a clear coating of the chromosomes by Esrrb but not by Nanog (Fig. 1C). We conclude, therefore, that global coating of the chromosomes can be captured using alternative fixatives to PFA, in particular bifunctional molecules that are expected to increase the speed and efficiency of cross-linking of protein-protein interactions within complexes.

**Figure 1:**
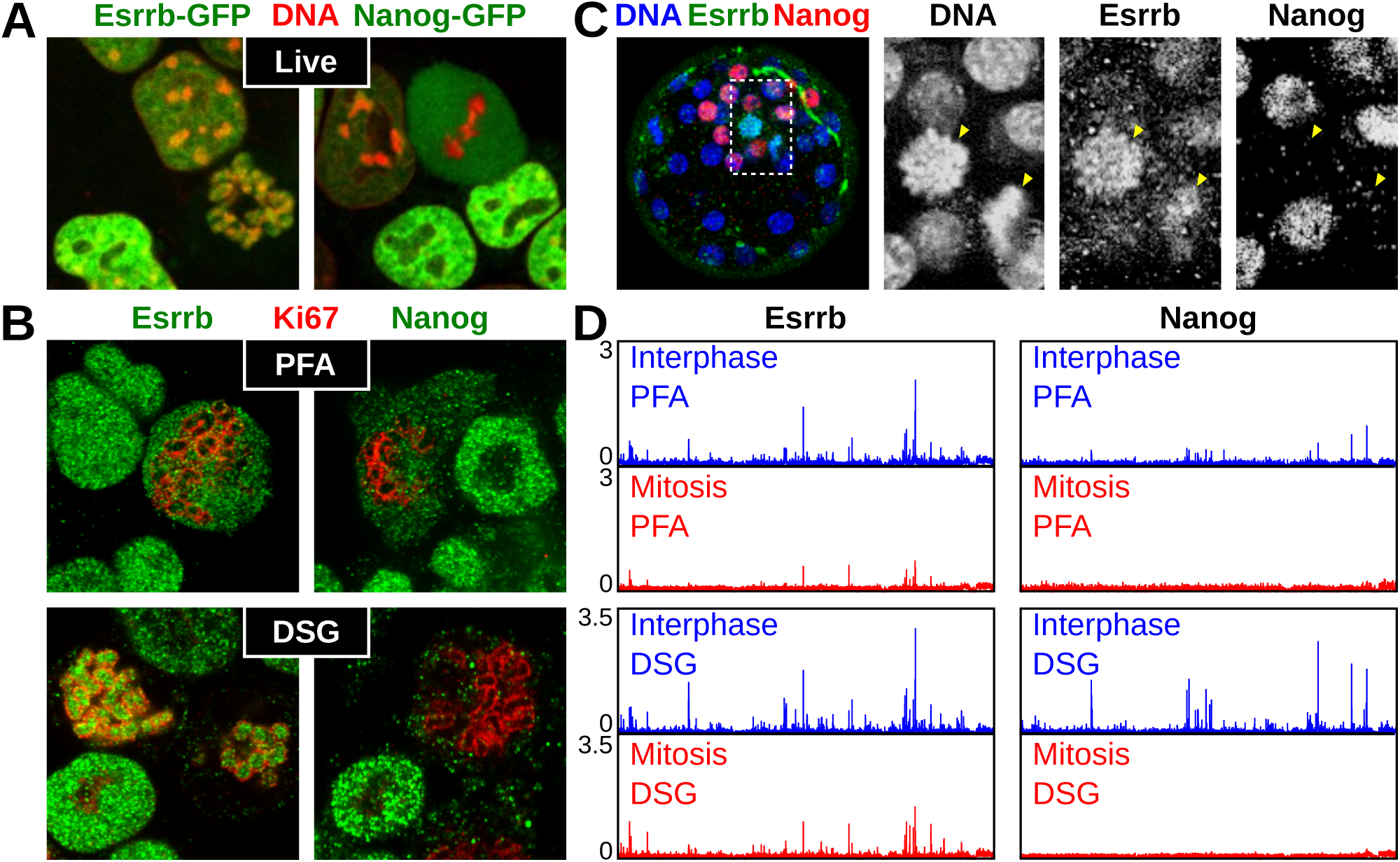
Capturing global Esrrb binding on mitotic chromosomes. **(A)** Localisation of Esrrb-GFP (left) or Nanog-GFP (right) fusion proteins in live cells cultured with Hoechst 33342 (red). **(B)** Esrrb (left) and Nanog (right) immunofluorescence (green) after fixation with either PFA (top) or DSG+PFA (bottom; annotated as DSG). The chromosome periphery of mitotic chromosomes is identified by Ki67 (red). **(C)** Immunostaining for Nanog (red) or Esrrb (green) performed on a mouse blastocyst. Counterstain with Hoechst 33342 is shown in blue. Close-up on two mitotic cells (arrowheads) is shown in the right panels (dashed area delimits the selected region). **(D)** Representative binding profiles of Esrrb and Nanog in reads per million (RPM) in interphase (blue) or mitosis (red), obtained after fixation with either PFA (top) or DSG+PFA (bottom; annotated as DSG). The region shown corresponds to chr17:25954686-27500000 (1.5 Mb)

### DSG fixation does not alter the profile of Esrrb binding in mitotic cells

Our finding that DSG and Glyoxal maintain the global association of Esrrb with the mitotic chromosomes opens the possibility to test whether this binding results from the sum of site-specific interactions or from other mechanisms. Indeed, if the global staining reflected site-specific interactions exclusively, one should expect to identify a much larger number of Esrrb binding sites by Chromatin Immuno-Precipitation (ChIP) after fixation with DSG or Glyoxal than with PFA. Yet, despite our efforts, we could not perform ChIP with these two reagents (data not shown). Since DSG or Glyoxal alone are sufficient in their own to globally retain Esrrb on mitotic chromosomes, this indicates that the underlying interactions are likely to be DNA-independent. After a dou-ble crosslinking with DSG followed by PFA (DSG+PFA), which is frequently used in biochemical approaches **(Tian et al. 2012)**, ChIP was instead particularly efficient (Figs. 1, 2, 4). Therefore, we performed ChIP-seq in asynchronous (thereafter interphase) and mitotic preparations of ES cells (>95% purity); after splitting the populations in two, we proceeded in parallel with either PFA or DSG+PFA crosslinking. We observed very similar profiles of Esrrb binding both in interphase and in mitosis, irrespective of whether the cells had been crosslinked with PFA or with DSG+PFA (Fig. 1D and Fig. 4). Therefore, whereas Esrrb is globally cross-linked outside or within the mitotic chromosomes by PFA and DSG respectively, the mitotic ChIP signal does not vary dramatically. We note however that DSG+PFA provides higher signal and better signal to background ratio, both in interphase and in mitosis. In agreement with immunostaining and live imaging, Nanog binding is globally lost, both in PFA and in DSG+PFA (Fig. 1D). From this analysis, we conclude that the global coating and the interaction of Esrrb with specific sites are two distinct phenomena. While mitotic Esrrb bookmarking (i.e. binding to specific sites) can be revealed by PFA and by DSG+PFA, global coating is only visible with DSG (or Glyoxal). If the former is a result of the DNA binding activity of Esrrb **(Festuccia et al. 2016)**, the molecular interactions underpinning the latter remain enigmatic. We can exclude, however, that DNA-independent Esrrb binding occurs exclusively at the periphery of the chromosomes, a proteinaceous compartment that includes TFs **(Booth and Earnshaw 2017)**. Indeed, Esrrb is detected covering the entire area delimited by Ki-67 (Fig. 1B and Fig. S1). This indicates the global Esrrb coating of the chromatids detected by microscopy reflects aspecific interactions of this TF with the chromatin.

**Figure 2:**
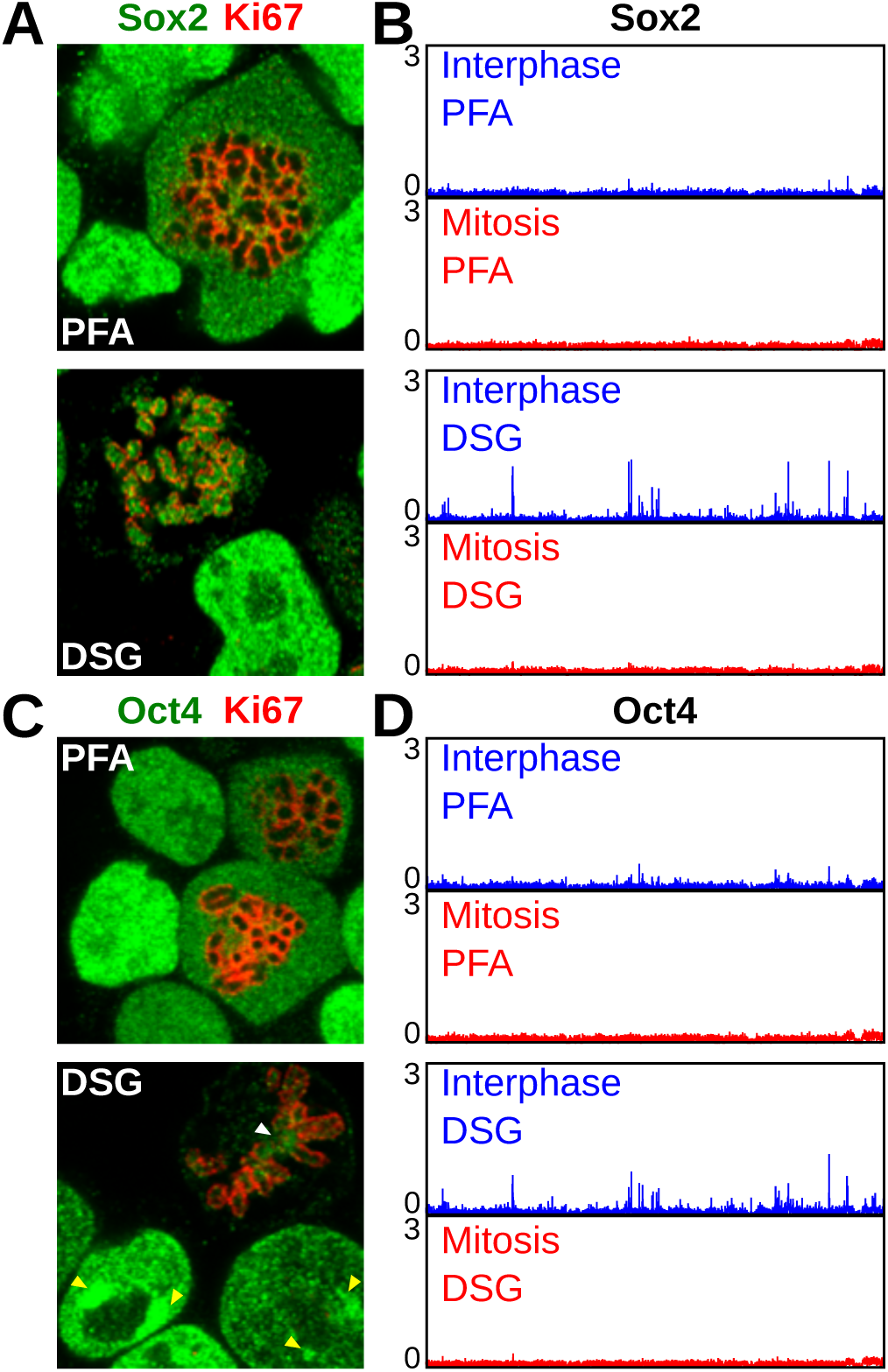
Sox2 and Oct4 do not bind at regulatory regions in mitosis. **(A)** Sox2 immunofluorescence (green), after fixation with either PFA (top) or DSG+PFA (bottom; annotated as DSG). The mitotic chromosome periphery is identified by Ki67 (red). **(B)** Representative binding profiles of Sox2 presented as in Fig. 1D. **(C, D)** Results of the same analyses described in (A) and (B) are shown for Oct4. Arrowheads indicate peri-centric heterochromatin foci (PCH) in interphase (yellow) and centromeres in mitosis (white).

**Figure 4:**
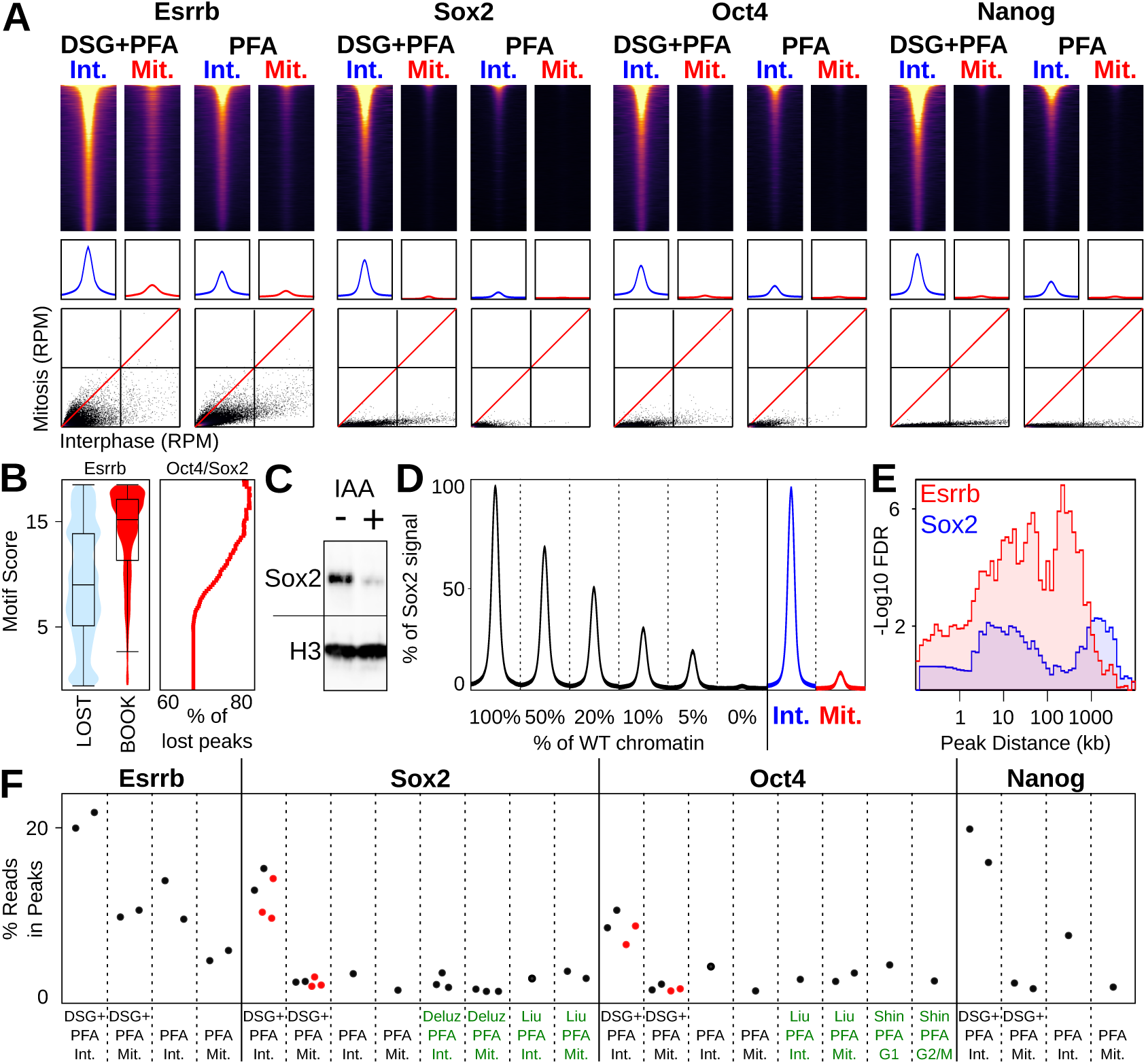
Comparative analysis of Esrrb, Sox2, Oct4 and Nanog binding in interphase and in mitosis. **(A)** Top: heatmaps of ChIP-seq signal at Esrrb, Sox2, Oct4 and Nanog binding regions identified in Interphase (Int.) and mitosis (Mit.) for DSG+PFA and PFA alone (see Methods for details). Middle: average binding profile of the regions shown in the heatmaps. Heatmaps and average binding profiles are scaled to the mean occupancy of each factor measured in interphase after DSG+PFA fixation. Bottom: scatter plots of ChIP signal (RPM) at above regions for interphase and mitosis (DSG+PFA scale 0-40 RPM; PFA scale 0-20 RPM). **(B)** Violin plots (left) depicting the FIMO-called best motif score per Esrrb peak in sites losing binding in mitosis (LOST) or retaining binding (BOOK). Right: percentage of Lost peaks with a composite Oct4/Sox2 motif of at least a given quality score. **(C)** Levels of Sox2-AID fusion protein in cells cultured in the absence (-) or presence (+) of the Auxin analogue IAA for 2 hours; H3 is shown as control. **(D)** Percent of the Sox2 ChIP signal detected at binding regions after spiking increasing amounts of WT chromatin into chromatin prepared from Sox2-depleted cells shown alongside the average Sox2 binding profile at potentially bookmarked regions in WT cells in interphase and mitosis. **(E)** Enrichment of genes responsive to Esrrb (red) and Sox2 (blue) in early G1 with proximity to Esrrb or Sox2 bookmarked regions, displayed as -log10 Fisher FDR for genes within given distance of a bookmarked peak. **(F)** Percentage of ChIP-seq reads in identified binding sites for Esrrb, Sox2, Oct4 and Nanog, in both interphase (Int.) and mitosis (Mit.) and DSG+PFA or PFA fixation in our data (black labels) and public datasets (green labels). The red dots correspond to the samples that were added to our study to further corroborate our results.

### DSG versus PFA comparisons reveal different behaviours of other proposed BFs

In addition to Esrrb, other pluripotency TFs have been proposed to act as BFs in ES cells **(Deluz et al. 2016; Teves et al. 2016; Liu et al. 2017)**, although evidence is contradictory. Sox2 has been consistently shown to globally associate with the chromosomes in three independent studies **(Deluz et al. 2016; Teves et al. 2016; Liu et al. 2017)**. In contrast, while one study **(Deluz et al. 2016)** reported that Sox2 binds with poor efficiency to a few dozen regions in mitosis (compared to thousands of sites in interphase), another study claimed that Sox2 and Oct4 remain bound to virtually all their interphase targets **(Liu et al. 2017)**. In addition, Sox2 and Oct4 were shown to be phosphorylated by Aurora kinases, which inhibits DNA binding in mitotic cells **(Qi et al. 2016; Shin et al. 2016)**. In these studies, ChIP was performed after PFA fixation, which leads to an apparent depletion from mitotic chromosomes of both Sox2 and Oct4 (Fig. 2A, C). In contrast, we found that Sox2 displays bright signal all over the chromosomal arms, within the Ki-67 delimited region, by immunostaining after DSG and Glyoxal fixation (Fig. 2A and Fig. S1). We thus extended our ChIP-seq analysis based on DSG+PFA fixation to Sox2. Whereas DSG+PFA dramatically increases ChIP efficiency of Sox2 compared to PFA, the profiles in mitosis are very similar for both fixatives, with little evidence for mitotic bookmarking activity (Fig. 2C and Fig. 4). Therefore, while displaying a macroscopic behaviour similar to Esrrb, mitotic Sox2 does not appear to be an efficient BF. Next, we analysed Oct4 binding. By immunofluorescence, we observed a nearly complete depletion from the chromosomal arms in mitosis, both after DSG and Glyoxal (Fig. 2B, C and Fig. S1). In agreement, ChIP-seq analysis also showed almost complete loss of Oct4 binding at its interphase targets (Fig. 2D). Our results are in agreement with a number of other studies **(Deluz et al. 2016; Shin et al. 2016; Kim et al. 2018)**. However, even using DSG+PFA we could not reproduce recent results showing mitotic bookmarking by Sox2 and Oct4 **(Liu et al. 2017)**. The use of inhibitors of MEK/GSK3b in the conflicting publication, which leads to a reinforcement of the pluripotency network’s activity, cannot account for these differences (Fig. S2). We conclude from our data that neither Sox2 nor Oct4 can be considered as potent BFs. Residual signal can be detected at some regions (Fig. 2B, Fig. 4 and Fig. S2), but not to levels comparable to Esrrb. In line with previous results, this further argues for the existence of two components driving the interaction of TFs with mitotic chromosomes: DNA-independent enrichment on the chromatin is detected for many, but not all, TFs; site-specific interactions with regulatory elements are a property of selected bookmarking factors like Esrrb.

### DSG enables capturing transient interactions at different chromatin compartments

Careful examination of the Oct4 stainings after DSG and Glyoxal, but not PFA fixation, unmasked a previously unnoticed accumulation of this TF at DAPI-rich regions, the chromocenters **(Saksouk et al. 2015)**, where several centromeres cluster together to form the pericentric heterochromatin (PCH; yellow arrowheads in Fig. 2C and in Fig. S1). Moreover, in mitotic cells we could also observe focal enrichment of Oct4 at centromeric regions (white arrowheads in Fig. 2C and in Fig. S1). This characteristic pattern of co-localisation with the PCH was further validated by live imaging using ectopic Oct4-GFP and endogenously expressed Oct4-RFP fusion proteins (Fig. 3A and Fig. S3A). Same results were obtained in cells cultured in regular conditions (Fig. 3A) or with inhibitors of MEK/GSK3b (Fig. S3A). In the latter conditions, PCH shifts from H3K9me3 to H3K27me3 **(Tosolini et al. 2018)**, indicating that the PCH association of Oct4 is independent of the presence of specific heterochromatic marks. Remarkably, Oct4 staining is similar to that of Aurora kinase b (Fig. S3B), which has been shown to phosphorylate Oct4 in mitotic cells to inhibit DNA binding. In agreement, in the presence of the Aurora kinase inhibitor Hesperadin a slight increase of Oct4 coating throughout the chromosomal arms could be observed (Fig.S3C). Hence, using alternative fixatives to PFA not only enables the visualisation of the genuine mitotic localisation of TFs, but may also reveal additional activities in interphase. We then asked whether the interactions of Sox2 and Oct4 unmasked by DSG and Glyoxal are indeed dynamic, as generally reported **(Teves et al. 2016)**. We observed highly dynamic interactions, both in interphase and in mitosis, for all three factors fused to GFP and analysed in parallel experiments (Fig. 3B, 3C, and Fig. S4). Esrrb and Oct4 displayed faster fluorescent recovery after photobleaching (FRAP) in mitosis (Fig. 3B, C). This is particularly true for the interaction of Oct4 with PCH, which are already very dynamic in interphase (Fig. 3C). In reciprocal experiments, we assessed fluorescence loss in photobleaching (FLIP; Fig. S4). We could not identify any significant remnant signal on mitotic chromatids after one minute of continuous bleaching of the freely diffusing TF molecules. Hence, DSG (and Glyoxal) are capable of capturing the highly dynamic interactions established by Esrrb/Sox2 on the chromosomal arms, and by Oct4 in PCH, both in interphase and in mitosis. Altogether, we conclude that while Esrrb exhibits robust mitotic book-marking activity, other factors are largely evicted during mitosis, irrespectively of their DNA-independent localisation to the chromatin.

**Figure 3:**
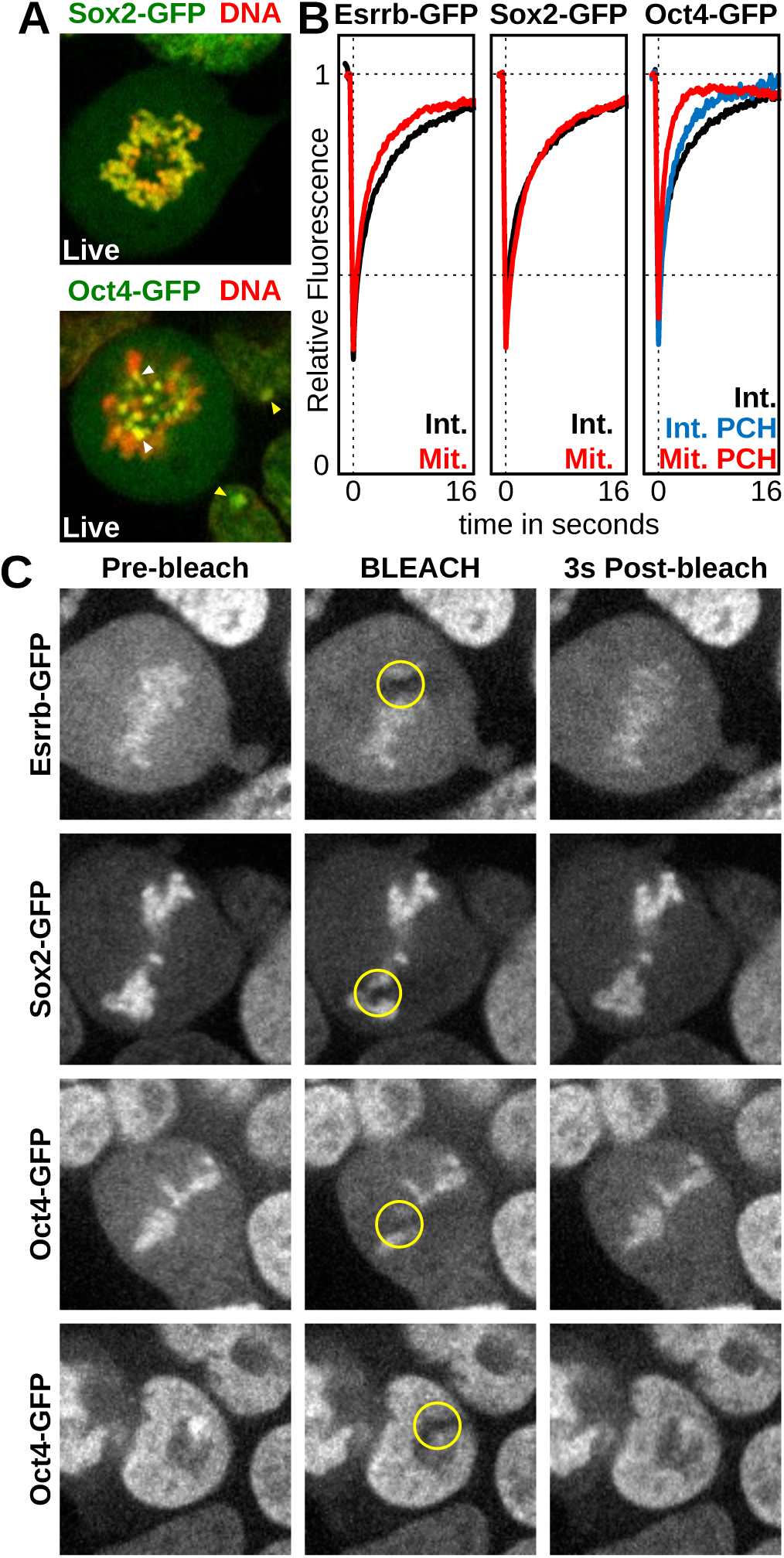
The interactions captured by DSG are dynamic. **(A)** Localisation of Sox2-GFP (top) or Oct4-GFP (bottom) fusion proteins (green) in live cells cultured with Hoechst 33342 (red). Arrowheads indicate peri-centric heterochromatin foci (PCH) in interphase (yellow) and centromeres in mitosis (white). **(B)** Quantifications of FRAP experiments in interphase (black) and mitosis (red) performed in cells expressing Esrrb-GFP or Sox2-GFP. For cells expressing Oct-4-GFP, recovery of fluorescence at (blue), or outside of (black), peri-centric heterochromatin foci (PCH) is displayed for interphase. Recovery at PCH is displayed for mitosis (red). The Y axis shows the mean percentage of fluorescence relative to pre-bleach levels detected in multiple independent experiments; the X axis shows the time after bleaching. **(C)** Representative examples of Esrrb-GFP, Sox2-GFP and Oct4-GFP signal on mitotic chromosomes before and after bleaching, at the indicated time (seconds). For Oct4-GFP the recovery of signal at PCH is also shown for cells in interphase (bottom).

### Esrrb is the only prominent BF among Esrrb, Sox2, Oct4 and Nanog

Using the collection of datasets generated for Esrrb, Sox2, Oct4 and Nanog in interphase and in mitosis, we sought to comprehensively identify regions subject to mitotic book-marking. To this end, we first identified the binding regions of individual TFs (Table S2) and confirmed that only Esrrb displays clear and frequent binding in mitosis (Fig. 4A); for Oct4, Sox2 and Nanog, only the regions displaying very high levels of binding in interphase show residual ChIP signal in mitosis, especially in DSG+PFA, where the number of detected peaks is increased. Peaks that were called only in DSG+PFA, and neither in our PFA samples nor in other publicly available datasets **(Chen et al. 2008; Marson et al. 2008; Aksoy et al. 2013; Whyte et al. 2013)**, tend to be smaller (Fig. S5). Nevertheless, their signal is clearly above background in all the analysed datasets of interphase cells fixed with PFA (Fig. S5). Hence, DSG helps capturing regions displaying low levels of binding and increases the overall efficiency of the ChIP. Nonetheless, it does not specifically unmask binding in mitosis. We then used a statistical differential occupancy approach to define regions as book-marked or lost (see Methods for details and Table S2). We found 10144 regions book-marked by Esrrb, representing 29.9% of its interphase sites. All other factors displayed a drastic contraction in binding in mitosis: 574 regions for Sox2 (2% of interphase targets); 102 regions for Oct4 (0.6%); 18 regions for Nanog (0.07%). Strong Esrrb binding motifs were identified at the vast majority of Esrrb book-marked regions (73.4%, score > 12), but only at a smaller subset of the regions losing binding in mitosis (34.9%, score > 12). In contrast, regions losing Esrrb binding displayed an increased occurrence of Oct4/Sox2 composite motifs (Fisher p<7e-45, Oct4/Sox2 Motif score > 12). Remarkably, we observed a scaling relationship: regions containing high quality Oct4/Sox2 motifs, exhibit a higher tendency to lose Esrrb binding in mitosis (Fig. 4B). Altogether, this indicates that at book-marked regions, Esrrb occupancy is primarily driven by specific interactions with the cognate binding sequence for this TF, as we have previously shown **(Festuccia et al. 2016)**. At lost regions, Esrrb is instead likely recruited indirectly by other TFs that are not capable of binding in mitosis. Hence, the substantial reduction of Esrrb binding sites observed in mitosis represents a striking example of the effect of loss of cooperative TF binding. Previously, we used titration experiments to investigate whether the binding levels seen for Esrrb in mitosis could be explained by contamination from interphase **(Festuccia et al.2016)**; all our mitotic preparation have less than 5% of remnant interphase cells, and typically between 2 and 4%. We repeated this analysis for Sox2, given the relatively high number of low mitotic peaks that we detected in comparison to Oct4 and Nanog. To generate Sox2depleted chromatin, we generated an ES cell line with (i) both endogenous Sox2 alleles tagged with an auxin-inducible degradation domain (Sox2-AID), and (ii) a constitutive transgene expressing the Tir1 protein inserted at the TIGRE locus **(Madisen et al. 2015)**. Upon treatment with the auxin analogue IAA for 2h, a significant reduction of Sox2 protein levels was observed (Fig. 4C). To further deplete Sox2, cells were differentiated in the presence of retinoic acid (RA) and IAA for 4 days. Gradually increasing amounts of WT chromatin were then spiked into chromatin prepared from IAARA treated cells, and ChIP-seq analysis performed. We found that as little of 5% of WT chromatin was sufficient to detect clear Sox2 peaks of reduced enrichment (Fig. 4D). Strikingly, the amount of signal observed by adding 5% of contaminant chromatin was higher, on average, to that seen in mitosis at the regions potentially book-marked by Sox2 (Fig. 4E). Therefore, it is possible that a significant fraction of the regions seemingly bound by Sox2 in mitosis, as well as the absolute levels of enrichment in mitosis, results from the small percentage of contaminant interphase cells in our preparations. To further corroborate that Sox2 is not an efficient book-marking factor we turned to a functional assay. Confirming our previously result, the set of Esrrb bookmarked regions identified here (Table S2) tend to be enriched in the vicinity of genes that are controlled by this TF in early G1 (Fig. 4E) **(Festuccia et al. 2016)**. We then introduced a GFP-Ccna cell-cycle reporter **(Festuccia et al. 2016)** into Sox2-AID cells, treated them with IAA for 2h and sorted early G1 cells to perform RNA-seq analyses. In comparison with Esrrb, we found a rather minor statistical association between the genes controlled by Sox2 in early G1 (Table S3) and the regions potentially bookmarked by Sox2 (Fig. 4E). We conclude that, whilst we cannot fully rule out that Sox2 may display minimal bookmarking activity, only Esrrb represents a potent and functionally relevant BF among the tested pluripotency factors. This conclusion is particularly well illustrated when the ChIP signal measured at each region is plotted in interphase versus mitosis (Fig. 4A; bottom panels), or when the proportion of reads on peaks are calculated for each TF (Fig. 4F). Why Sox2 and Oct4 have been previously found mitotically bound at most of their interphase targets **(Liu et al. 2017)** remains therefore unclear. This is particularly striking, taking into consideration that our DSG+PFA datasets clearly display improved ChIP efficiency compared to several other published profiles (Fig. 4F; Deluz et al. 2016; Shin et al. 2016; Liu et al. 2017). Despite our efforts, and the addition of 3 and 2 additional independent replicates for Sox2 and Oct4, respectively (red dots in Fig. 4F), we did not find strong evidence for Sox2 and Oct4 bookmarking.

### Drastic changes in nucleosome organisation characterise regulatory elements in mitosis

Recently, mitotic chromatin has been shown to maintain surprisingly high levels of chromatin accessibility at virtually all regulatory elements that are active in interphase, in particular at promoters **(Hsiung et al. 2015; Teves et al. 2016)**. Accordingly, we observed that promoter accessibility in mitotic ES cells even surpasses the level observed in interphase, as evaluated by ATAC-seq (Fig. 5A). However, distinct nucleosome organisations might characterise accessible chromatin in these two phases of the cell cycle **(Kelly et al. 2010; Rhee et al. 2014; Mieczkowski et al. 2016; Voong et al. 2016; Mueller et al. 2017)**. To address this, we inferred nucleosome positioning and stability in interphase and in mitosis from a series of experiments based on MNase-seq and H3 ChIP-seq using chromatin digested with titrated MNase activity. In mitosis, we observed preserved nucleosome depleted regions (NDR) around the transcription start sites of promoters (TSS; Fig. 5B). Yet, the phasing of nucleosomes at both sides of the NDRs was drastically attenuated in mitotic cells, probably reflecting reduced transcriptional activity (Fig. 5B). Moreover, when we compared average H3 ChIP-seq signal between mitosis and interphase at different levels of MNase digestion (Fig. 5C), a clear asymmetry was revealed: upstream of the TSS, the sensitivity of the nucleosomes to MNase increased in mitotic cells (as shown by reduced signal with strong digestion); downstream, the +1 nucleosome displayed a similar stability than in interphase, while the following nucleosomes acquired in mitosis increasing levels of fragility. At the minimal promoter region (TSS and 150bp upstream), we did not find evidence of a nucleosome displaying high occupancy either in interphase or in mitosis (Fig. 5B). Nonetheless, the H3 signal detected over the minimal promoter tend to increase in mitosis, irrespectively of the MNase conditions (Fig. 5C). These results indicate that, globally, the nucleosomes at promoters are more fragile **(Mieczkowski et al. 2016; Voong et al. 2016)** in mitosis, except at the minimal promoter region where they stabilise without increasing their overall occupancy. Moreover, the differential behaviour within and outside the transcription unit may potentially reflect the reduced transcriptional activity that has been recently detected in mitotic cells **(Palozola et al. 2017)**. Therefore, promoters are subject to drastic nucleosome reorganisation in mitotic cells. We then analysed enhancers (identified here as p300bound elements, excluding TSSs and gene bodies). As previously shown **(Hsiung et al. 2015)**, we found enhancers to partially lose accessibility in mitosis (Fig. 5D). More strik-ingly, these elements display a profound reconfiguration in nucleosomal architecture (Fig. 5E): nucleosomes resistant to our most aggressive digestion conditions can be detected at the site of p300 recruitment exclusively in mitosis, and the phasing of the surrounding nucleosomes is altered (Fig. 5E). Moreover, titration of MNase activity followed by H3 ChIP-seq, revealed that both upstream and downstream of the stabilised nucleosome, increased fragility can be measured in mitotic cells (Fig. 5F). Therefore, even though promoters and enhancers maintain significant levels of accessibility in mitotic cells, the arrangement of their nucleosomes changes substantially.

**Figure 5:**
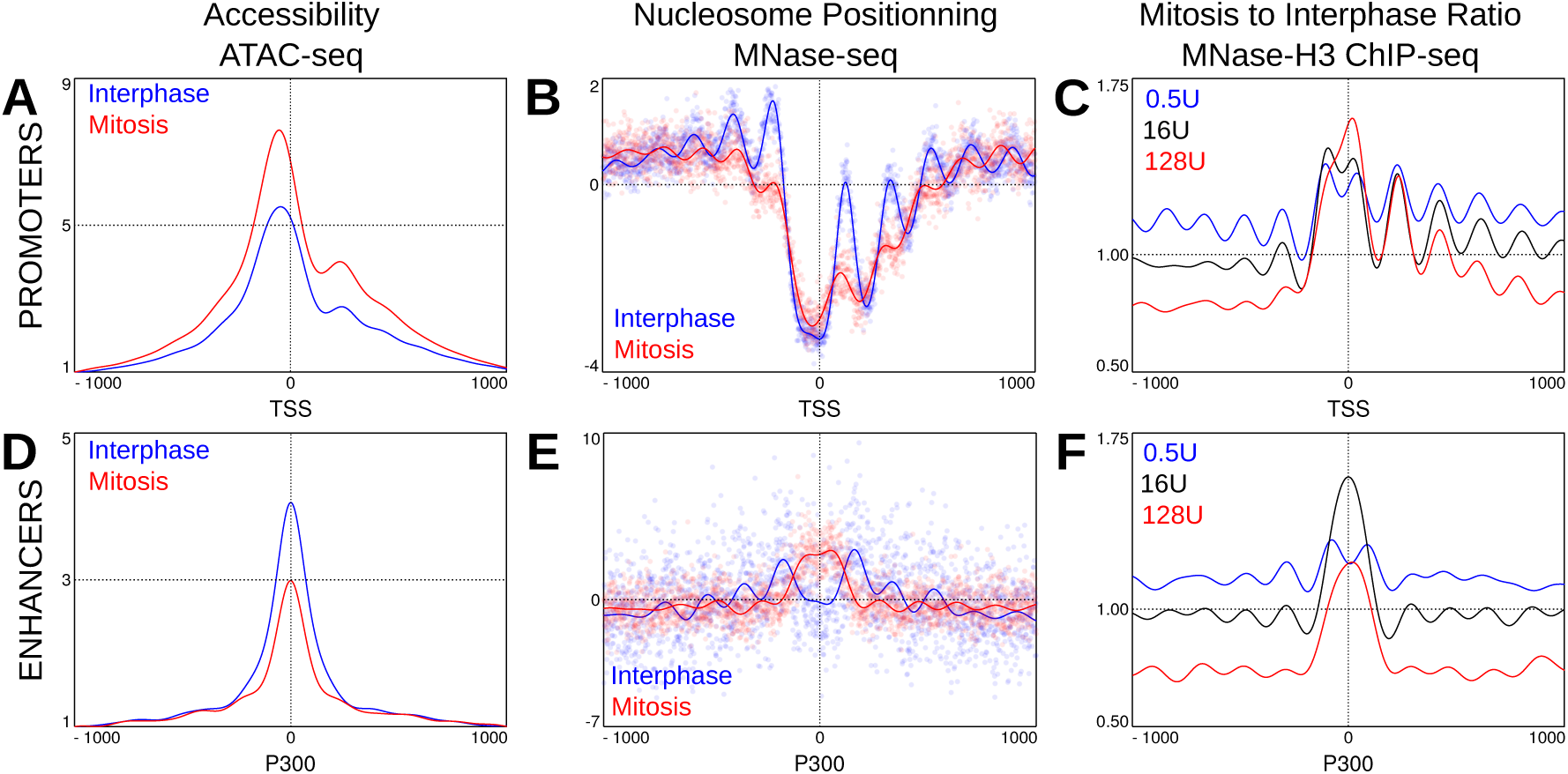
Drastic reconfigurations of promoters and enhancers in mitotic cells. **(A)** Accessibility profiles measured by ATAC-seq in the region surrounding the TSS of active genes in interphase (blue) or mitosis (red). Signal is number of Tn5 cut sites for 0-100 bp fragments, normalised to minimum accessibility in +/1000 bp window. **(B)** Nucleosome positioning at the same set of promoters, established by MNase-seq. In this panel, the z-score of the number of midpoints of nucleosome-sized fragments (140-200bp) per base, after digestion with 16U of enzyme, are plotted. The lines represent a Gaussian process modelling nucleosome positioning (see Methods) in interphase (blue) and in mitosis (red). **(C)** Mitosis over interphase ratio of MNase H3 ChIP-seq signal for nucleosomal fragments (as assessed by Gaussian process regression; see Methods). Ratios shown for MNase digestions with 0.5U (blue), 16U (black) and 128U (red) of enzyme. **(D, E, F)** as (A, B, C) but for regions centred on summits of interphase P300 ChIP-seq peaks excluding promoters. Note that in (E) MNase-seq signal is from 128U digestions.

### Chromatin accessibility and nucleosome organisation as a function of Esrrb book-marking

We then focused on the analysis of the regions bound by Esrrb (Fig. 6 and Fig. S6). While Esrrb-book-marked regions partially lose accessibility (Fig. 6A), this reduction is significantly more pronounced at the regions where Esrrb binding is lost in mitosis (Fig. 6C). Hence, there is a clear correlation between the ability of Esrrb to bind to certain targets in mitotic cells, and the partial maintenance of accessibility. Moreover, at book-marked regions, we observed highly positioned nucleosomes both in interphase and mitosis: the Esrrb motif lies within a major NDR and phased nucleosomes spread both upstream and downstream the binding site (Fig. 6B). This pattern contrasts markedly with that seen at p300 enhancers (Fig. 5E), clearly establishing a strong correlation between Esrrb mitotic binding and the retention of well-structured nucleosome arrays. Moreover, in mitosis we observed a slight shrinking of the nucleosomal array converging towards the central Esrrb motif, leading to a modest change of position of the nucleosomes. Remarkably, when we calculated a frequency map of additional Esrrb motifs within these regions (grey histogram in Fig. 6B), we observed a small but clear enrichment precisely at the mitosis-specific inter-nucleosomal space between the −2/−1 and +1/+2 nucleosomes. This strongly indicates that in mitosis, the DNA binding activity of Esrrb becomes dominant in establishing nucleosome positioning. In contrast, at regions losing Esrrb binding in mitosis, the nucleosomal profiles were not found to be dramatically different in interphase and mitosis: in both cell-cycle phases the Esrrb motif is occupied by a nucleosome, which is more sharply positioned during division (Fig. 6D). However, these regions appeared barely organised compared to their book-marked counterparts and lacked clear phasing at both sides of the Esrrb motif. Since high quality Esrrb motifs are not particularly prevalent at these regions (Fig. 4B), we reanalysed the data by re-centring on Esrrb summits. We noted that Oct4/Sox2 motifs are enriched in the vicinity of Esrrb summits (grey histogram in Fig. 6E), and therefore also recentred these regions on these motifs (Fig. 6F). Both analyses unveiled a clear nucleosomal organisation in interphase that is highly modified in mitotic cells (Fig. 6E, F). This indicates that Esrrb may be recruited indirectly and play a minor role in establishing nucleosome positioning over these regions. In accord, the nucleosome pattern at regions centred on Esrrb summits was also highly similar to that seen at the bulk of Oct4/Sox2 binding sites (Fig. 6H). These regions show a consistent reduction in accessibility in mitosis (Fig. 6G) and major nucleosome repositioning, with signs of shifting in the nucleosomal array and invasion at both flanks of the Oct4/Sox2 motifs (Fig. 6H). At all these regions, a concomitant increase in occupancy by fragile nucleosomes could also be observed (Fig. S6A). Of note, local features like the ones we observed at TSSs and p300 summits could not be detected (Fig. 5E, F and Fig. S6A). Finally, at regions exhibiting low mitotic Sox2 ChIP-seq signal, we also observed major reorganisations of nucleosomes in mitosis. Nonetheless, the presence of a very narrow NDR could not be ruled out (Fig. S6B), possibly reflecting minimal book-marking activity. From these analyses, we conclude that TF binding is likely required to maintain nucleosome positioning at regulatory elements during cell division. Esrrb acts as a major organiser of the chromatin in both phases of the cell cycle (Fig. 6B).

**Figure 6:**
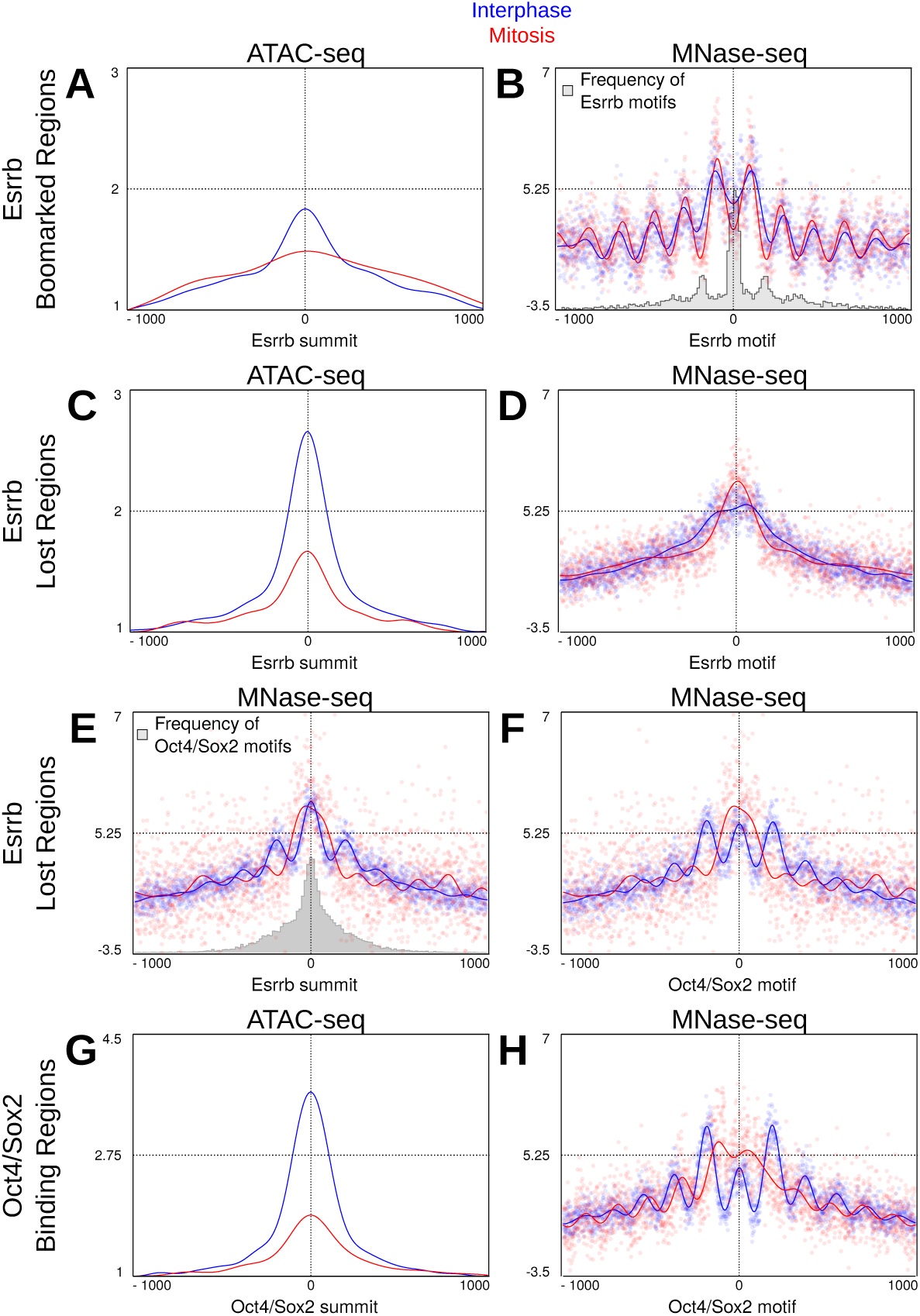
Binding of Esrrb at its cognate motif drives nucleosome organisation in interphase and in mitosis. Accessibility **(A, C, G)** determined by ATAC-seq as in Fig. 5A, and nucleosome positioning **(B, D, E, F, H)** established as in Fig. 5B, at the regions indicated on the left and centred as shown on their corresponding X-axis, in interphase (blue) and mitosis (red). Histograms embedded in (B) and (E), depict rate of occurrence of the indicated binding motifs: (B) additional Esrrb motifs with FIMO score > 8; (E) top scoring Oct4/Sox2 composite motif.

## Discussion

Proposed around 20 years ago **(Michelotti et al. 1997)**, the idea that certain TFs mitotically propagate gene regulatory information had been until recently only sporadically explored. Instead, over the last few years, several publications have revealed a continuously growing number of candidate mitotic book-marking TFs **(Festuccia et al. 2017)**. Considering that PFA, arguably the most used cross-linker, leads to an artificial depletion of TFs from the mitotic chromosomes, as visualised by microscopy, many more TFs than those currently described are probably able to associate with the chromatin during division **(Pallier et al. 2003; Teves et al. 2016)**. However, whether all these TFs are engaged in site-specific interactions and therefore act as mitotic book-marking factors remains unclear **(Festuccia et al. 2017)**. Here, we identify cross-linkers that preserve the global mitotic localisation of several TFs, providing a simple experimental method to study the behaviour of new transcriptional regulators during division and, more generally, visualise spatial organisations deriving from transient and fast binding events. Conversely, our results impose caution: we show that localisation of a TF to the chromatin does not necessarily imply sequence specific binding in mitosis (Fig. 7). This is exemplified by Sox2 and, as shown by others, by CTCF **(Oomen et al. 2018)**: while these TFs are both macroscopically retained, they are largely evicted from the sites occupied in interphase. The functional consequences of this distinction are major: we failed to identify a strong relationship between the proximity of the few regions exhibiting Sox2 binding, albeit at low levels, and the transcriptional effects of Sox2 in early G1. Conversely, the functional relevance of site-specific mitotic binding (Fig. 7) has been documented for several canonical book-marking factors, including Gata1 **(Kadauke et al. 2012)**, FoxA1 **(Caravaca et al. 2013)** and Esrrb **(Festuccia et al. 2016)**. Therefore, the emerging idea of a widespread mitotic book-marking activity needs to be carefully considered and evaluated. At the same time, the potential function of a global chromosomal retention cannot be ignored and requires dedicated experimental setups. In this regard, our comparative analysis of fixatives reveals that distinct molecular mechanisms likely contribute to the overall mitotic localisation of TFs (Fig. 7). Esrrb displays highly correlated binding profiles by ChIP when the chromatin is fixed with PFA or with DSG. In contrast, only DSG captures global Esrrb enrichment on the chromatin. Given the ability of DSG to efficiently fix transient interactions, and in light of the results of FRAP and single molecule tracking studies **(Caravaca et al. 2013; Deluz et al. 2016; Teves et al. 2016)**, this reveals that most likely the bulk of the molecules for a given TF bound to the chromatids during mitosis are not engaged in sequence specific interactions with DNA. Somehow surprisingly, however, we showed previously that mutating 3 aminoacids of the Esrrb DNA biding domain that are engaged in base-specific contacts with the binding motif dramatically decreases the global decoration of the mitotic chromosomes **(Festuccia et al. 2016)**. It is possible that these amino acids of the Esrrb zinc-finger domain are also required for Esrrb to scan the DNA in search of its biding sites. Alternatively, these mutations may more generally alter the structure of Esrrb, preventing interactions with other proteins enriched at the mitotic chromosomes. Notably, the bifunctional crosslinkers that we have used, DSG and Glyoxal, are expected to increase the efficiency of fixation within large protein complexes, opening the possibility that the interactions driving the global enrichment of TFs on the chromatids are based on protein-protein rather than protein-DNA contacts. Thus, we propose that the model previously proposed for FoxA1 regarding the existence of at least two distinct phenomena underlying the behaviour of TFs in mitotic cells could be extended, and applied generally to BFs: on the one hand, both DNA scanning and the ability to interact with other proteins of the chromatin sustains the bulk localisation of TFs to the chromatids; on the other, bona-fide book-marking, understood here as the capacity to mediate site-specific binding, drives functionally relevant accumulation of TFs at regulatory elements **(Festuccia et al. 2017)**. While FoxA1 is capable of binding nucleosomes directly **(Cirillo et al. 1998)**, by virtue of its inherent structural properties **(Clark et al. 1993; Ramakrishnan et al. 1993)**, the mitotic partners for the protein-protein interaction of other TFs decorating mitotic chromosomes may be more diverse (Fig. 7). These protein could be part of the chromatin or restricted to the chromosomal periphery **(Booth and Earnshaw 2017)**. While such restricted localisation can be excluded for Esrrb, Sox2 and Oct4, it may apply to other TFs. Indeed, a multitude of determinant of TF localisation seem to exist. This is the case of Oct4, that we report here as focally enriched within (peri-)centric regions, both in interphase and in mitosis. Extending beyond mitosis, given the complexity revealed by the used of multiple cross-linking agents, this study directly calls for a general reassessment of TF localisation and function as inferred from fixed samples.

**Figure 7:**
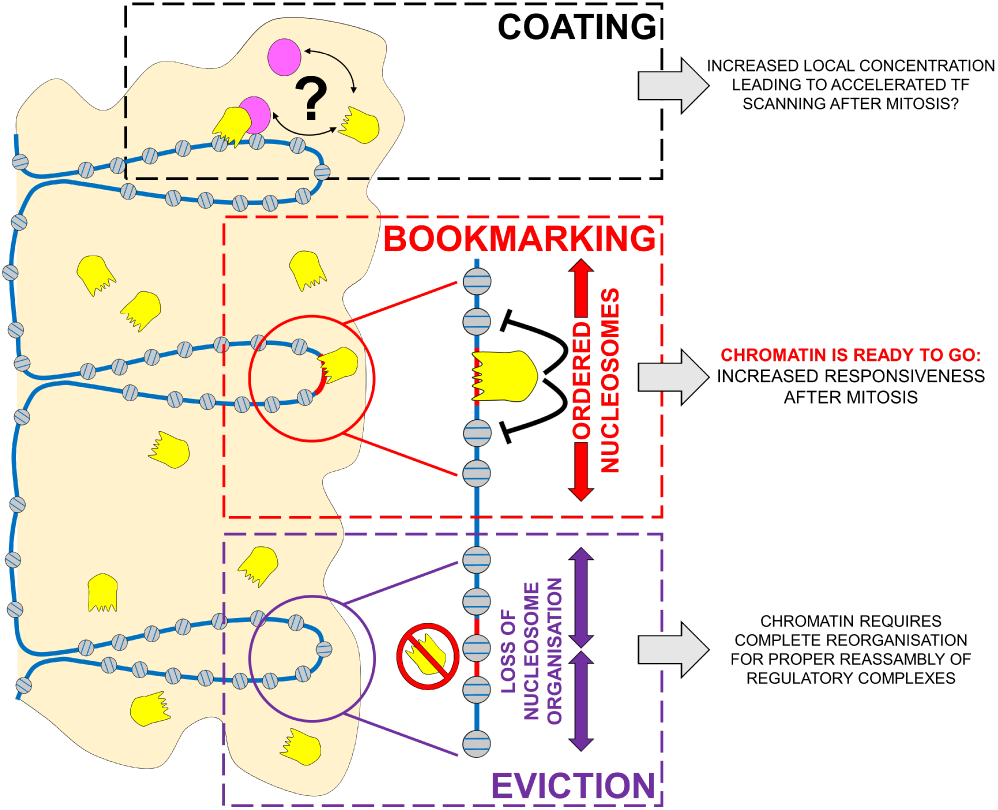
Model summarising distinct behaviours of TFs in mitotic cells and their relationships to nucleosome organisation and post-mitotic gene regulation. Many TFs show global localisation on the chromosomes in mitosis. This localisation is likely driven by sequence-independent interactions with DNA or other components of the chromatin or the mitotic chromosomes, and might serve a function in increasing the local concentration of TFs in proximity of their targets, in turn facilitating binding in G1. In contrast, during division only few TFs remain dynamically bound to a subset of the sites they occupy in interphase. At book-marked sites, the continued activity of these TFs maintains an ordered chromatin configuration, possibly limiting the extent of chromatin remodelling required to re-establish functional regulatory architectures in the following cell-cycle. At sites losing TF binding, nucleosome positioning is vastly disorganised, and increased occupancy by nucleosomes is detected at binding motifs. Although these sites do not become fully inaccessible, profound chromatin rearrangements are expected to be needed in early G1 to reinstate proper function.

Distinguishing TFs as enriched or depleted from mitotic chromosomes, and as binding or not at specific regulatory regions, will eventually allow us to establish a hierarchy of their contributions to the re-establishment of transcription after mitosis (Fig. 7). This will be particularly important in highly proliferative cells undergoing progressive implementation of new cell identities during development **(Festuccia et al. 2017)**. To gain a full understanding of the importance of mitotic book-marking, it is also crucial to elucidate the molecular mechanisms mediating its function. Different lines of evidence point to the lack of permanent TF binding at mitotic chromosomes. Even in the extreme case of the general TF Tbp, the residence time on the mitotic chromatin is below 2 minutes **(Teves et al. 2018)**. Therefore, occupancy by single molecules of mitotic book-marking factors do not physically transfer regulatory information from mother to daughter cells; to be functional, BFs may instead induce specific modifications around their mitotic target sites. However, regardless of their mitotic book-marking status, most if not all active regulatory regions remain at least partially accessible in mitotic cells **(Hsiung et al. 2015; Teves et al. 2016)**. This has been now shown analysing the book-marking sites of several TFs, including Gata1 **(Kadauke et al. 2012)**, and, here, Esrrb, Sox2, Oct4 and Nanog. Therefore, even if many other BFs remain to be identified, the general loss of TF binding characterising mitosis is unlikely to completely abolish chromatin accessibility. In general, the presence of destabilised nucleosomes at regulatory elements could suffice to maintain these regions less refractory to the binding of transcriptional regulators. Nevertheless, TF binding might still contribute towards maintaining comparatively high accessibility at selected loci. This was originally proposed for the book-marking factor Foxl1 **(Yan et al. 2006)** and is further supported by our observation that the regions book-marked by Esrrb display a milder reduction of ATAC signal compared to those where Esrrb is evicted. More strikingly, our nucleosome mapping studies indicate that Esrrb book-marking plays a major role in preserving the fine patterns of nucleosome organisation, rather than mere accessibility, at regulatory elements (Fig. 7). Indeed, at regions bookmarked by Esrrb, binding motifs are strongly associated with a nucleosome depleted region, and are flanked by well organised and phased nucleosomes. This configuration is detected in interphase, but is significantly clearer in mitosis where even neighbouring inter-nucleosomal spaces correlate with the presence of additional Esrrb motifs. We believe this reflects the loss of counteracting effect from binding of other TFs in mitosis, and the consequent dominance of Esrrb over the organisation of the nucleosomes at these sites. In this light, mitosis might represent a context of simplified interactions of TF with the chromatin, where few fundamental activities are maintained. In contrast, in the complete absence of mitotic TF binding, nucleosomal arrays are largely reconfigured. This is true at enhancers marked by p300, at regions losing Oct4/Sox2 binding, as well as at CTCF biding sites **(Oomen et al. 2018)**. Remarkably, at regions losing Esrrb in mitosis a clear nucleosomal organisation is only appreciated when regions are aligned relative to the Esrrb peak summit or the binding motifs for other TFs. Hence, at these regions, Esrrb might be recruited indirectly and the nucleosomal organisation of these regions, therefore, is not imposed by Esrrb. Together these observations clearly indicate that mitotic bookmarking by Esrrb is essentially driven by sequence-specific DNA interactions through which this factor imposes specific constraints on nucleosomal organisation. Therefore, the nucleosomal landscape around TF binding sites in mitosis may be used as a proxy for mitotic bookmarking activity, further indicating that neither Sox2 nor Oct4 are efficient bookmarking factors.

The recent observation of widespread chromatin accessibility in mitotic cells suggested that many TFs would act as bookmarking factors. In contrast, our analysis of TF binding, chromatin accessibility, nucleosome positioning and stability in mitotic ES cells, rather indicates that mitotic bookmarking can only be mediated by selected TFs, such as Esrrb in ES cells. Indeed, the stereotypical behaviour of enhancers that we observe here indicates that a robust nucleosome is positioned at p300 recruitment sites, with more fragile nucleosomes occupying the vicinities. These destabilised nucleosomes may explain the apparent accessibility of these regions. At promoters, we also observe a loss of phasing, and a relative stabilisation of the nucleosomes lying just upstream of the TSS as compared to those more distally located, which appear to be more fragile. While the molecular players destabilising these nucleosomes requires further investigation, our data indicate that Esrrb, and potentially other bookmarking factors, may generally act by locally preserving specific nucleosome architectures. These configurations in turn favour the re-establishment of functional regulatory complexes early after mitosis. We propose this mechanism to represent the molecular basis of the transmission of regulatory information by sequence-specific mitotic bookmarking factors (Fig. 7).

## Acknowledgements

The authors acknowledge the Imagopole France–BioImaging infrastructure, supported by the French National Research Agency (ANR 10-INSB-04-01, Investments for the Future), for advice and access to the UltraVIEW VOX system. We also acknowledge the Transcriptome and EpiGenome, BioMics, Center for Innovation and Technological Research of the Institut Pasteur for NGS. This work was supported by recurrent funding from the Institut Pasteur, the CNRS, and Revive (Investissement d’Avenir; ANR-10-LABX-73). P.N. acknowledges financial support from the Fondation Schlumberger (FRM FSER 2017), the Agence Nationale de la Recherche (ANR 16 CE12 0004 01 MITMAT), and the Ligue contre le Cancer (LNCC EL2018 NAVARRO). N.F was supported by a Marie Curie IEF fellowship and a Pasteur-Cantarini Fellowship program. N.O. is supported by Revive.

## Author contributions

N.F. performed or supervised all the experimental work. N.O. analysed sequencing data. N.F., N.O. and P.N. analysed and interpreted the results and wrote the manuscript. All other authors provided experimental help.

## Declaration of interests

The authors declare no competing interests.

## SUPPLEMENTARY INFORMATION

Six supplementary figures accompany this manuscript:

**Figure S1:**
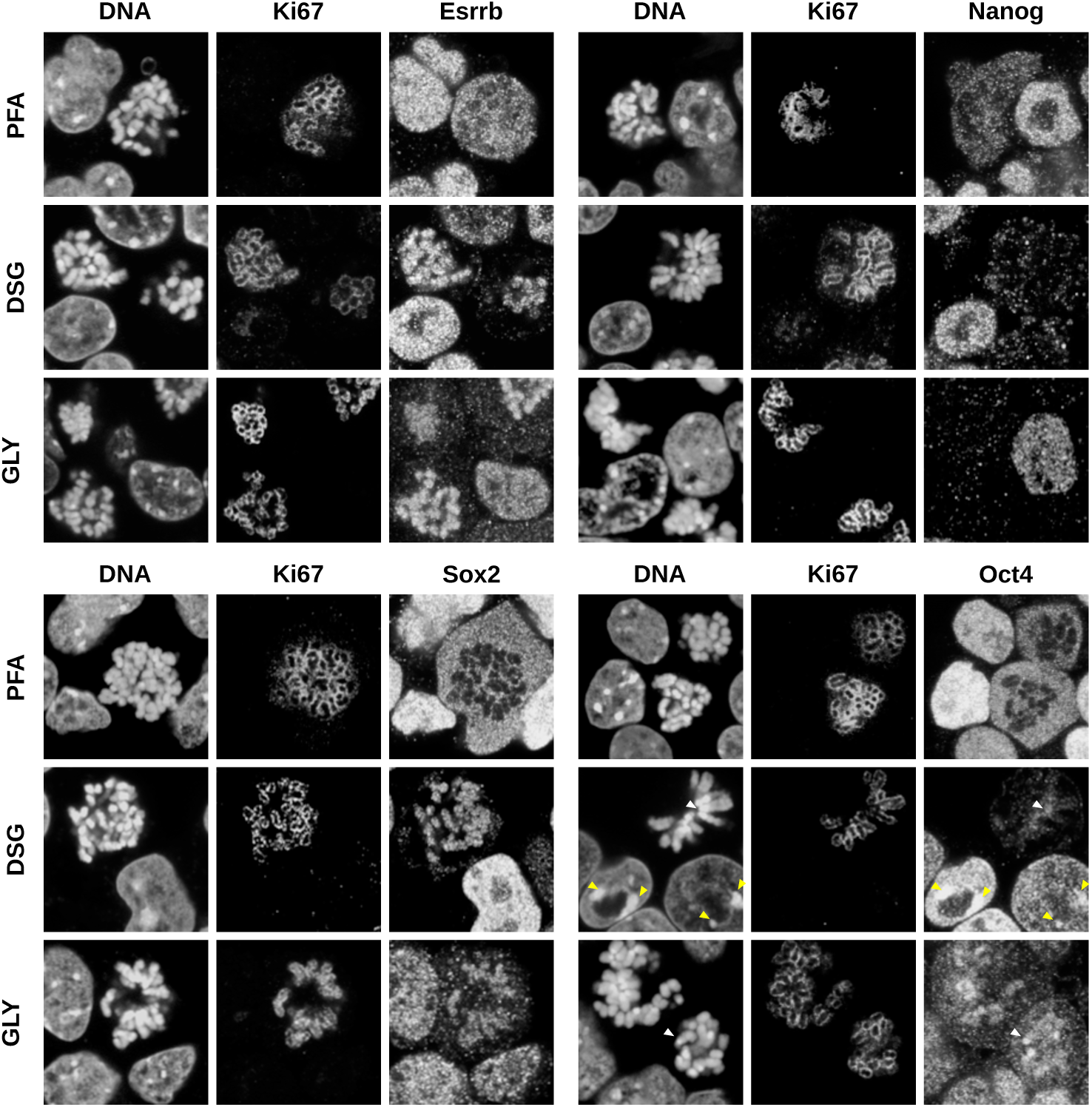
Comparative analyses of different fixations. Immunofluorescence for Esrrb (top left), Nanog (top right), Sox2 (bottom left) and Oct4 (bottom right), after fixation with either PFA, DSG+PFA (labelled as DSG only), or glyoxal. DNA was counterstained with DAPI. The mitotic chromosome periphery is identified by Ki67 staining. In the Oct4 staining, the arrowheads indicate peri-centric heterochromatin foci (PCH) in interphase (yellow) and centromeres in mitosis (white). Note that Sox2 immunofluorescene required a PFA post-fixation after Glyoxal.

**Figure S2:**
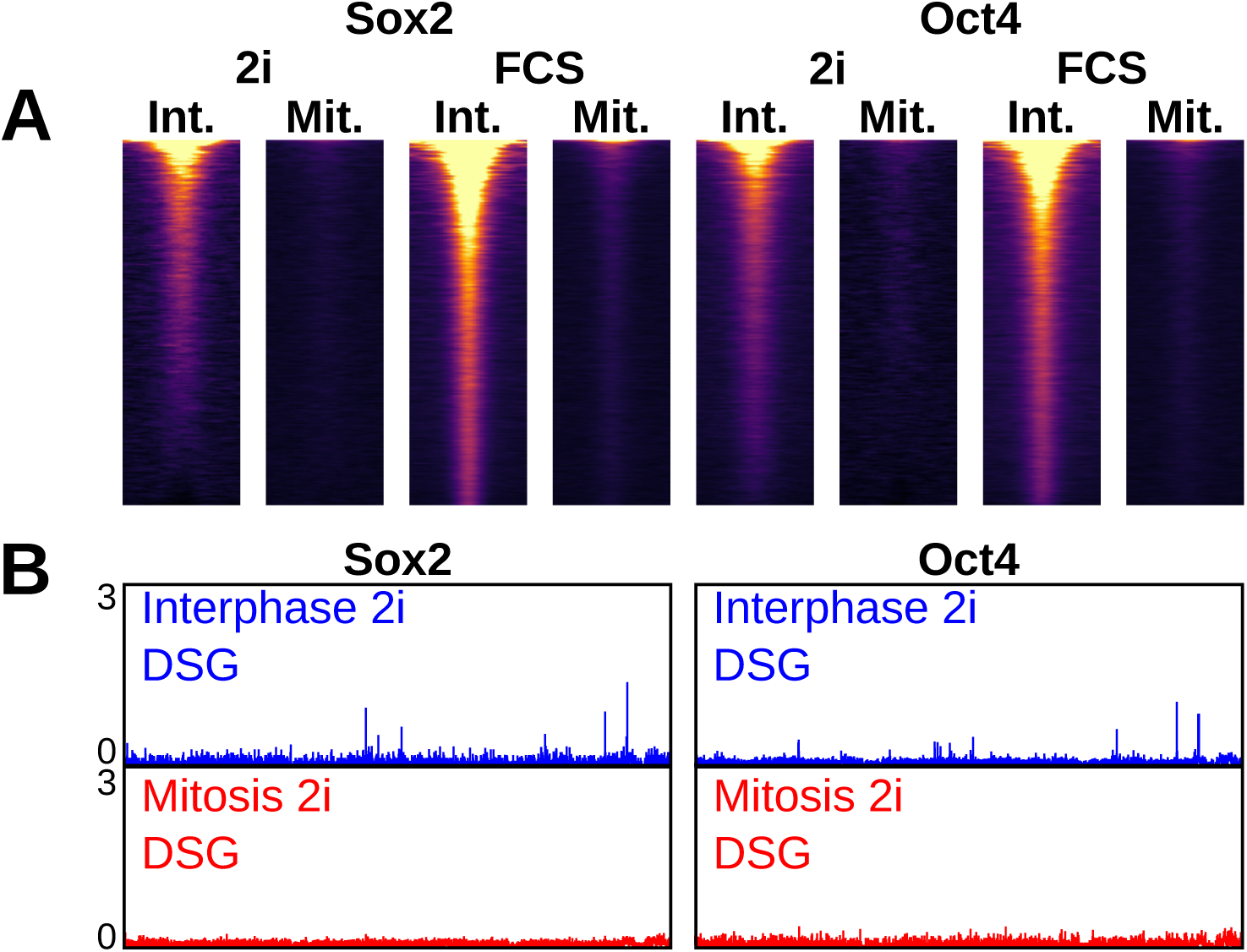
Oct4 and Sox2 binding in 2i-treated ES cells. **(A)** Heatmaps of ChIP-seq signal in interphase and mitosis for Sox2 and Oct4, in cells cultured in FCS/LIF with and without 2i. Binding regions are the union of peaks identified in both conditions. **(B)** Representative binding profile for Sox2 (left) or Oct4 (right) in interphase (blue) or mitosis (red), obtained after fixation with DSG+PFA in 2i treated cells; vertical scale RPM. The region corresponds exactly to that shown in Figs. 1 and 2.

**Figure S3:**
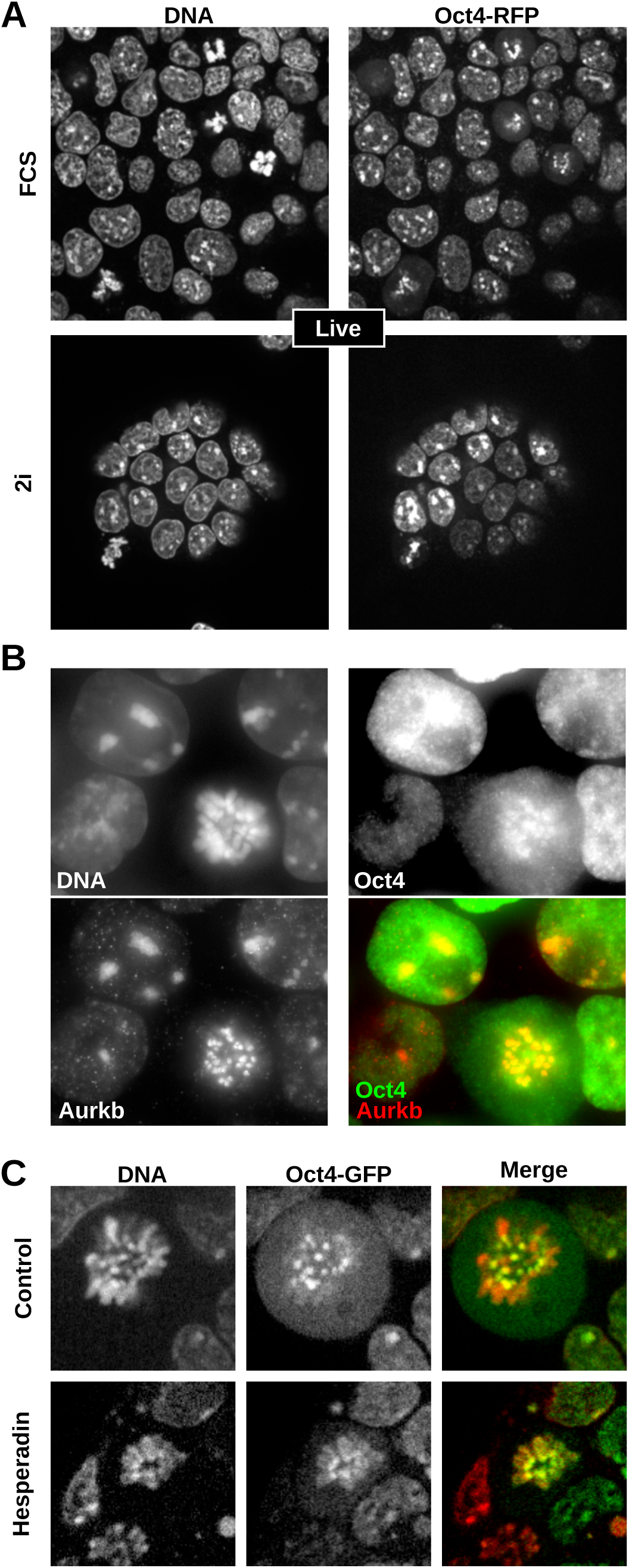
Extended analysis of Oct4 localisation. **(A)** Localisation of Oct4-RFP fusion proteins expressed from one of the two endogenous Oct4 alleles in live cells cultured in FCS/LIF medium (top) on in FCS-free 2i-containing medium (bottom). DNA is visualised by Hoechst 33342 (red). **(B)** Oct4 (green in the merge) and Aurkb (red in the merge) immunofluorescence after fixation with DSG. Note the large overlap at PCH and at centromeres in interphase and in mitosis. **(C)** Localisation of Oct4-GFP fusion proteins in live cells cultured in FCS/LIF medium supplemented (bottom) or not (top) with the Aurkb inhibitor Hesperadin. DNA is visualised by Hoechst 33342 (red). Note this image corresponds exactly to that shown in Fig. 3A.

**Figure S4:**
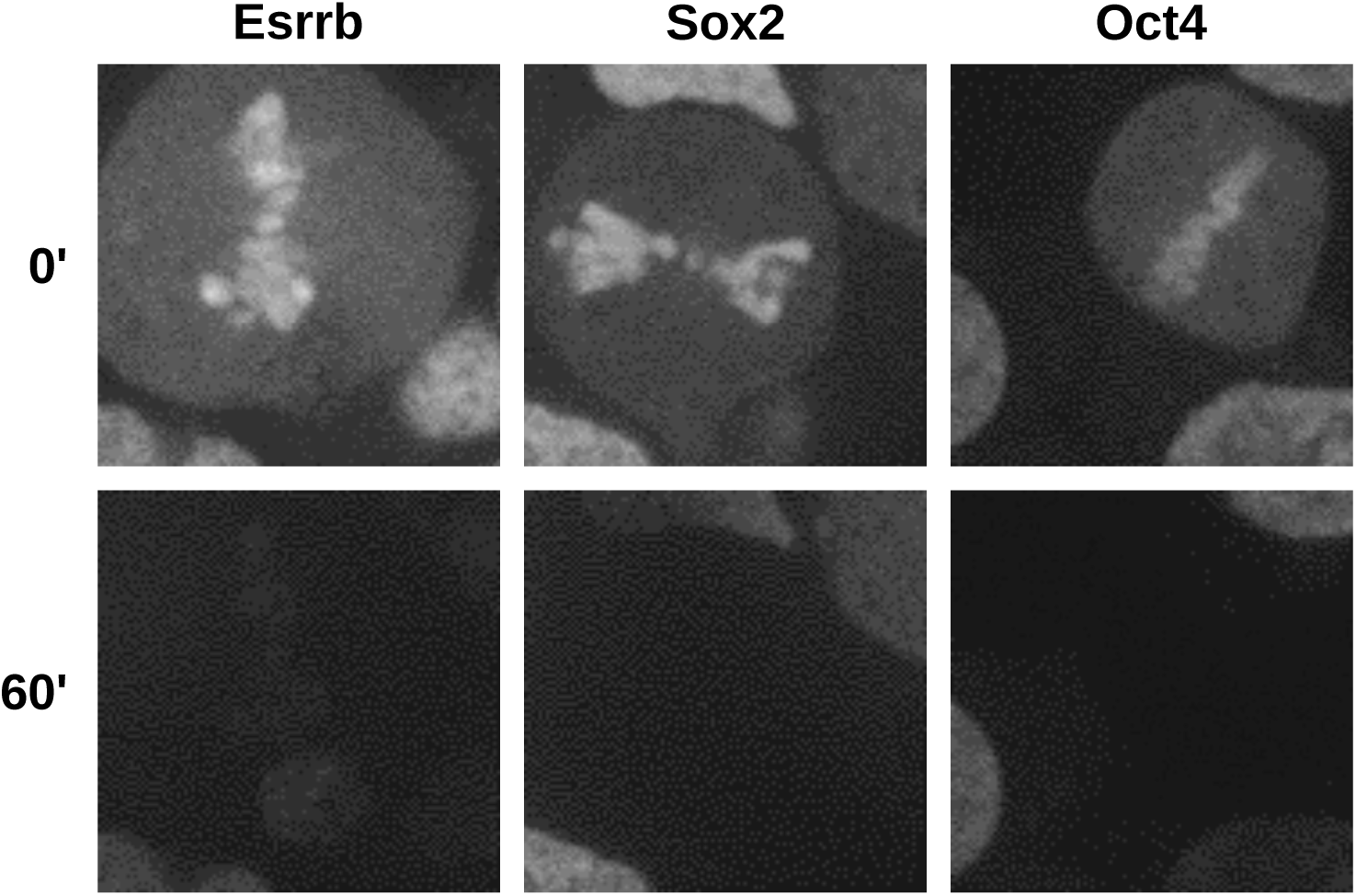
Example of FLIP imaging. Representative examples of Esrrb-GFP, Sox2-GFP and Oct4-GFP signal on mitotic chromosomes before and after 60 second of continuous bleaching of freely diffusing molecules outside the chromatids.

**Figure S5:**
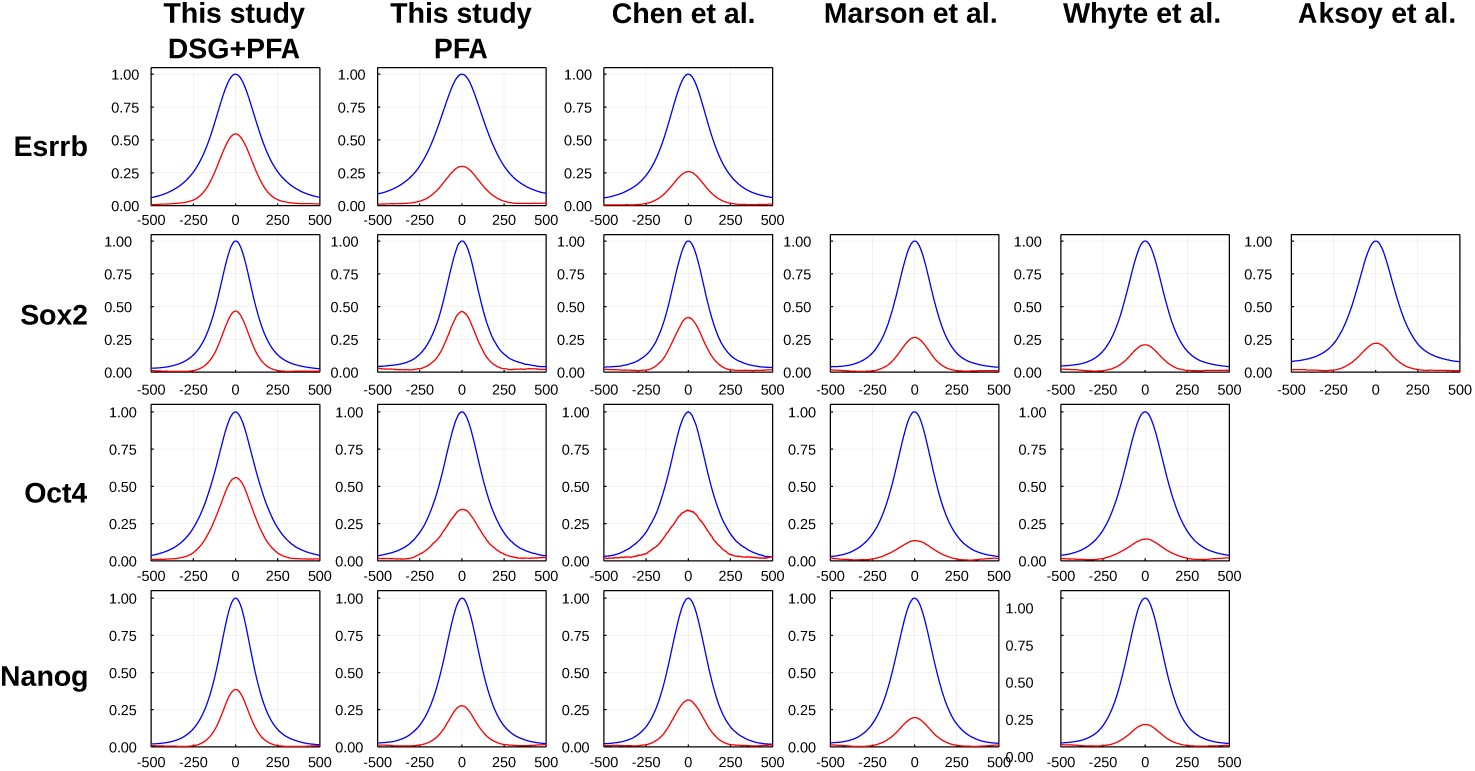
Average binding profile in interphase of Esrrb, Sox2, Oct4 and Nanog, in this study and in public datasets. Blue line depicts all binding regions identified in this study, red depicts regions detected in DSG+PFA exclusively, i.e. regions with no significant peak in any of the indicated PFA dataset.

**Figure S6:**
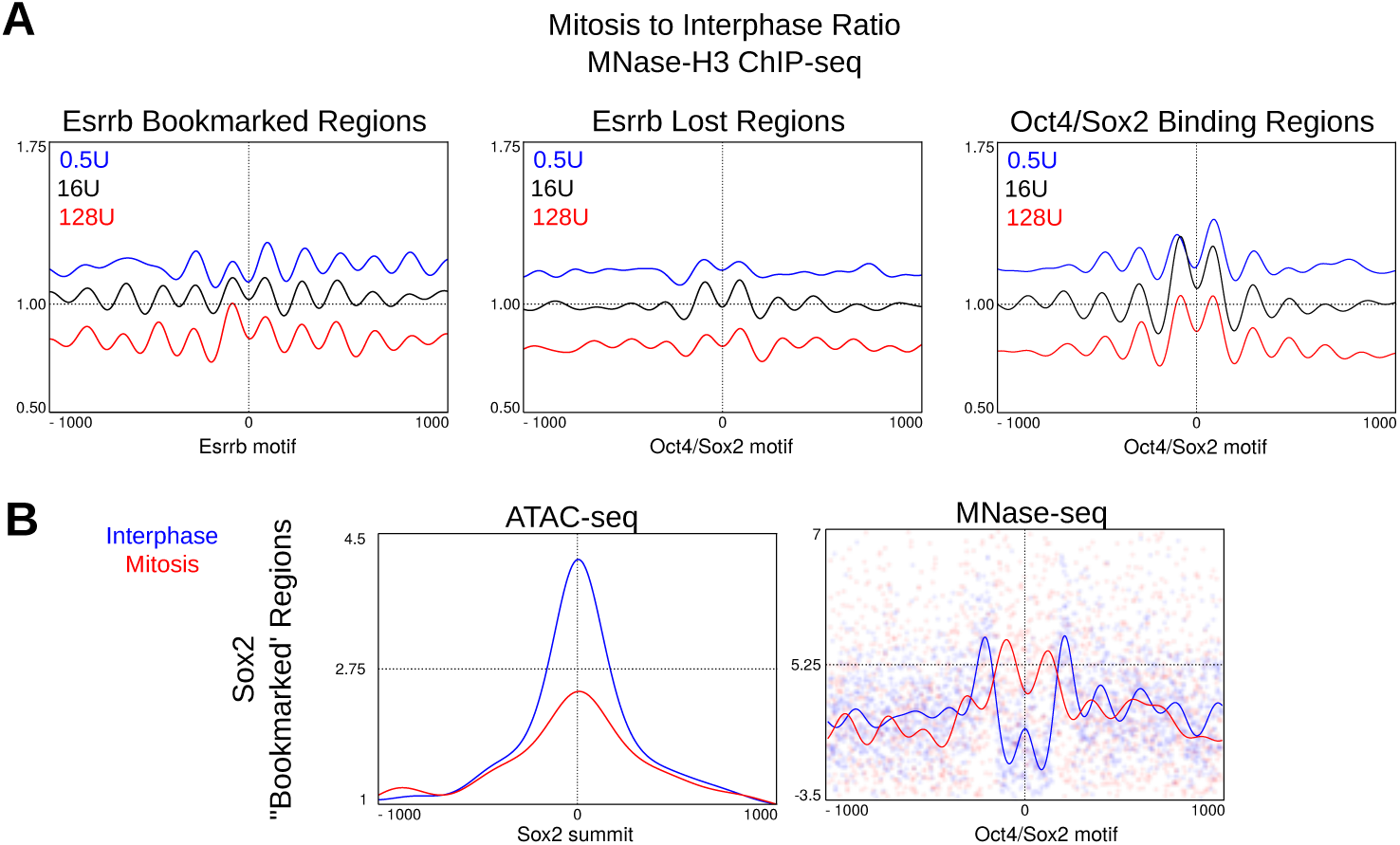
Additional information of nucleosome organisation at Esrrb, Oct4 and Sox2 sites. **(A)** Ratio of MNase H3 ChIP-seq nucleosomal size fragment signal of mitosis over interphase, as described in Fig 5, for 0.5U (blue), 16U (black) and 128U (red) MNase concentrations, at Esrrb bookmarked and lost regions and all Oct4+Sox2 binding regions, centred on the top Esrrb motif, top Oct4/Sox2 motif and top Oct4/Sox2 motif respectively. **(B)** Left: Accessibility measured by cut sites of 0-100 bp ATAC-seq fragments at Sox2 putative bookmarked sites in interphase and mitosis, centred on Sox2 peak summits. Right: Nucleosome positioning measured by MNase-seq nucleosomal size fragments (140-200 bp) after digestion with 16U at Sox2 putative bookmarked sites centred on Oct4/Sox2 motif. Vertical scale gives z-score.

They can be found at the end of this document.

Three Supplementary Tables are available online:

**Table S1:** Overview of ChIP-seq, MNase-Seq, MNase H3 ChIP-seq, ATAC-seq, RNA-seq libraries sequenced in this study.

**Table S2:** Peaks and bookmarking calls for Esrrb, Oct4, Sox2, Nanog

**Table S3:** Differential expression of genes responsive to Sox2 depletion in EG1.

